# Velocity of Conduction Between Columns and Layers in Barrel Cortex Reported by Parvalbumin Interneurons

**DOI:** 10.1101/2022.07.27.501767

**Authors:** Kate S. Scheuer, John Judge, Xinyu Zhao, Meyer B. Jackson

## Abstract

Inhibitory interneurons that express parvalbumin (PV) play critical roles throughout the brain. Their rapid-spiking characteristics enable them to control the dynamics of neural circuits across a range of time scales, but the timing of their activation by different cortical pathways remains unclear. Here, we use a genetically encoded hybrid voltage sensor, hVOS, to image PV interneuron voltage changes with sub-millisecond precision in primary somatosensory barrel cortex (BC) of adult male and female mice. Electrical stimulation evoked depolarizing responses with a latency that increased with distance from the stimulating electrode, allowing us to determine conduction velocity. By focusing on conduction between cortical layers or between barrel columns we were able to measure interlaminar or intralaminar conduction velocity, respectively. Velocities ranged from 74 to 473 μm/msec depending on trajectory, and we found that interlaminar conduction velocity was about 71% faster than intralaminar conduction velocity. This suggests that computations within columns can be processed more rapidly than between columns. BC circuitry integrates thalamic and intracortical input for a variety of functions including texture discrimination and sensory tuning. The difference in timing between intra- and interlaminar activation of PV interneurons could impact these functions. This study illustrates how hVOS imaging of PV interneuron electrical activity can reveal differences in the dynamics of signaling within different elements of cortical circuitry, and this approach offers a unique opportunity to investigate conduction in populations of axons based on their targeting specificity.

## Significance Statement

Parvalbumin (PV) interneurons play critical roles throughout the brain, shaping sensory responses with their distinct properties and circuitry. Here we use the genetically encoded hybrid voltage sensor, hVOS, to measure conduction velocity along excitatory axons targeting PV interneurons in the primary somatosensory barrel cortex (BC). We find that conduction velocity within a barrel column is 71% faster than between columns. This suggests that the BC prioritizes PV interneuron-related computations within columns. Functionally, these differences between interlaminar and intralaminar conduction velocity could impact processes including texture discrimination, sensory tuning, and integration of thalamic and cortical inputs.

## Introduction

Expression of the protein parvalbumin (PV) defines a broad class of inhibitory interneurons which have been implicated in psychiatric and neurological disorders including schizophrenia, bipolar disorder, and autism spectrum disorder (Liu et al., 2012; Gonzalez-Burgos et al., 2015; Lauber et al., 2018). Because PV interneurons fire rapidly and have a short membrane time constant, they play a critical role in controlling the temporal integration window of their targets (Pouille and Scanziani, 2001; Galarreta and Hestrin, 2002; Cardin, 2018; Ferguson and Gao, 2018). PV interneurons are distributed throughout the cortex, including the whisker-related primary somatosensory cortex, or barrel cortex (BC), where their anatomy, circuitry, and function have received much attention (Staiger and Petersen, 2021). BC is divided into visually distinct cytoarchitectural units, or barrels, giving the region its name (Woolsey and Van der Loos, 1970). Each barrel receives input from one vibrissa. Barrel column boundaries, visible in L4, can be extended through supragranular and infragranular layers (Lefort et al., 2009; Staiger and Petersen, 2021). Whisker inputs follow the canonical circuit, passing through the lemniscal pathway from the ventral posteromedial thalamus (VPM) to a distinct barrel in L4 before being relayed to L2/3 and then L5. However, BC also contains a number of non-canonical pathways, including those extending between two or more columns, and receives long-range inputs from a variety of areas including multiple thalamic regions, primary motor, auditory, and visual cortices, and secondary motor and somatosensory cortices (Staiger and Petersen, 2021). Functionally, BC has been implicated not only in touch and whisking (Feldmeyer, 2012; Staiger and Petersen, 2021) but also in sensing movement (Ayaz et al., 2019), texture discrimination (Allitt et al., 2017; Isett et al., 2018; Vecchia et al., 2020), object localization (O’Connor et al., 2010), and social interactions (Lenschow and Brecht, 2015). However, how BC processes and integrates spatial and temporal sensory input is not clear.

PV interneurons are present in L2-6 in BC and participate in essentially all BC responses, enhancing spatial and temporal resolution, enforcing sparse coding, and providing both feedback and feedforward inhibition (Staiger and Petersen, 2021; Yeganeh et al., 2022). PV interneurons also contribute to computations across columns and cortical layers. Excitation/inhibition balance and gamma oscillation frequency, both linked to PV interneuron function, differ between cortical layers (Xu et al., 2016; Adesnik, 2018). The fast-spiking nature of PV interneurons suggests that they are adapted to rapid computations in which the speed of their activation is likely to be important to their functions. Measurement of conductance velocity in the activation of PV interneurons across cortical layers and columns is needed to address this hypothesis.

Here we use the genetically encoded hybrid voltage sensor hVOS to compare the timing of activation of PV interneurons between cortical layers within a column (interlaminar), as well as within a layer between columns (intralaminar). Imaging reveals responses from many cells distributed through multiple cortical layers and columns with sub-millisecond temporal resolution (Chanda et al., 2005; Wang et al., 2010; Ghitani et al., 2015). By targeting hVOS probe to PV interneurons (Bayguinov et al., 2017), we were able to track the spread of their activation by excitatory axons that run within and between layers. Our measurements of velocity generally fell in the range of previously reported values in cortex. We found that interlaminar conduction velocity within a column is 71% faster than intralaminar conduction velocity across columns. Thus, PV interneuron-related computations within columns can be faster than those between columns, and this is likely to have functional consequences within and beyond BC.

## Materials and Methods

### Animals

Ai35-hVOS1.5 (C57BL/6-*Gt(ROSA)26Sor^tm1(CAG-hVOS1.5)Mbja^*/J, https://www.jax.org/strain/031102) Cre reporter mice (Bayguinov et al., 2017) were bred with PV Cre driver mice (B6.129P2-Pvalb^tm1(cre)Arbr^/J, https://www.jax.org/strain/017320) to create animals with PV interneuron-specific hVOS probe expression. All animal procedures were approved by the Animal Care and Use Committee of the University of Wisconsin-Madison School of Medicine and Public Health (IACUC protocol: M005952).

### Hybrid voltage sensor (hVOS)

The hVOS probe used here harbors a cerulean fluorescent protein (CeFP) tethered to the inner cell membrane by a truncated h-ras motif (Wang et al., 2010). Slices are perfused with 4 μM dipicrylamine (DPA), a small anion which partitions into the cell membrane. Membrane depolarization drives DPA towards the CeFP to quench fluorescence through Förster resonance energy transfer, and repolarization drives the DPA away allowing fluorescence to return to baseline (Chanda et al., 2005; Wang et al., 2010). Fluorescence thus reports voltage changes selectively from PV interneurons because the PV Cre driver used here drives probe expression specifically and efficiently in these cells (Bayguinov et al., 2017). DPA has a response time < 0.5 msec (Chanda et al., 2005; Bradley et al., 2009), enabling hVOS fluorescence to track action potentials in single cells with excellent temporal fidelity (Ghitani et al., 2015; Ma et al., 2019).

### Slice preparation

Six- to ten-week-old male and female mice were deeply anesthetized with isoflurane and sacrificed via cervical dislocation. Brains were dissected and placed into ice-cold cutting solution (in mM: 10 glucose, 125 NaCl, 4 KCl, 1.25 NaH_2_PO_4_, 26 NaHCO_3_, 6 MgSO_4_, 1 CaCl_2_) bubbled with 95% O_2_/5% CO_2_. Coronal slices (300 μm) were prepared using a Leica VT1200S vibratome and placed into artificial cerebrospinal fluid (ACSF, same as cutting solution except 1.3 mM MgSO_4_, 2.5 mM CaCl_2_,) and bubbled with 95% O_2_/5% CO_2_ for at least one hour. ACSF used for slice storage and recording contained 4 μM DPA.

### Voltage imaging

Slices were continuously perfused with ACSF at room temperature and viewed using a BX51 Olympus microscope. Layer and barrel boundaries were visually identified in fluorescence and gradient contrast images (e.g., Fig 1A-F). Stimulus pulses (100 μA, 180-μsec, except Fig. 2A-C which used 10-60 μA) were generated by triggering a stimulus isolator (World Precision Instruments, Sarasota, Florida) and applied via fire-polished, ACSF-filled KG-33 glass electrodes (King Precision Glass, Claremont, California) with tip diameter ∼6-8 μm. Stimulus electrodes were positioned in various locations within BC with a micromanipulator. Displayed traces of fluorescence versus time were averages of 5 trials, at 15 sec intervals. Slices were illuminated with an LED light source (Prizmatix, Holon, Israel) with peak emission at 435 nm. Gradient contrast images were acquired with a high-resolution Kiralux CMOS camera (Thorlabs, Newton, New Jersey); fluorescence images were acquired with this camera for better visualization, but for voltage imaging we used a CCD-SMQ camera (RedShirt Imaging, Decatur, Georgia) at a framerate of 2000 Hz and 80×80 spatial resolution. Imaging was controlled with a custom acquisition and analysis program that gates stimulation and illumination (Chang, 2006).

**Figure 1.**
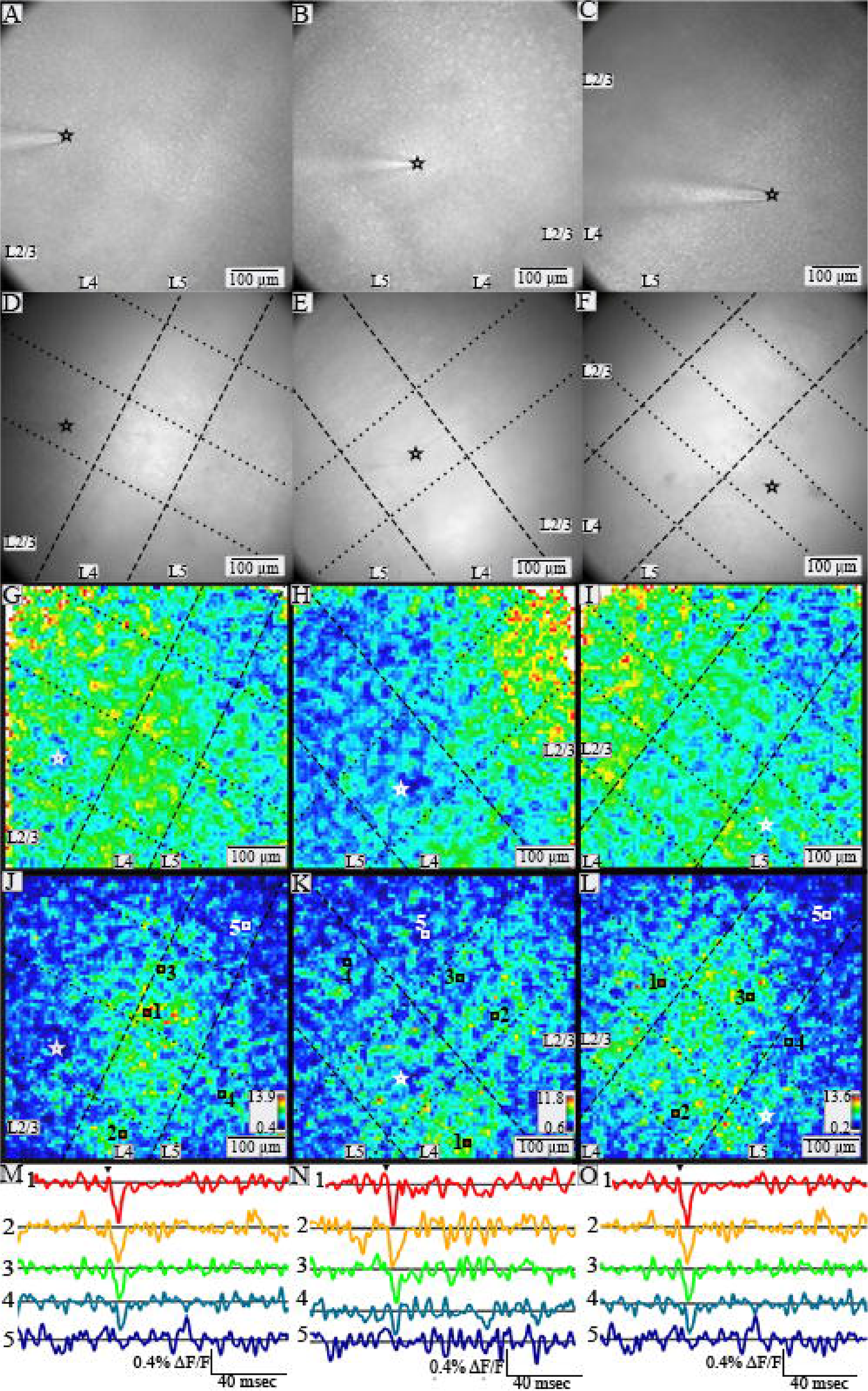
PV interneurons throughout a slice respond with different spatial patterns to stimulation in different layers. A-C. Gradient contrast images of slices of BC, all with L2/3 through L5 within the field of view. The tip of the stimulating electrode is visible in L2/3 (A), L4 (B), and L5 (C), and its position is indicated by black or white stars in panels A-L. D-F. Fluorescence images for the same slices in A-C. In D-L dashed lines show layer boundaries, and dotted lines show column boundaries. G-I. Maximum amplitude heatmaps for the corresponding slices above. Warmer colors correspond to larger peak amplitudes. A few white pixels at the edges reflect saturation due higher noise with lower light levels. J-L. SNR heatmaps for the corresponding slices above. Both maximum amplitude and SNR heatmaps are based on images acquired using different cameras than for gradient contrast and fluorescent images, so the fields of view do not align precisely. Warmer colors indicate higher SNR (scales and ranges indicated in lower right corners). Locations with higher SNR have stimulus-evoked responses in selected traces of fluorescence versus time (M-O) from numbered locations (indicated with white or black squares) in the corresponding SNR heatmaps above. Arrowheads above top traces indicate time of stimulation.

**Figure 2.**
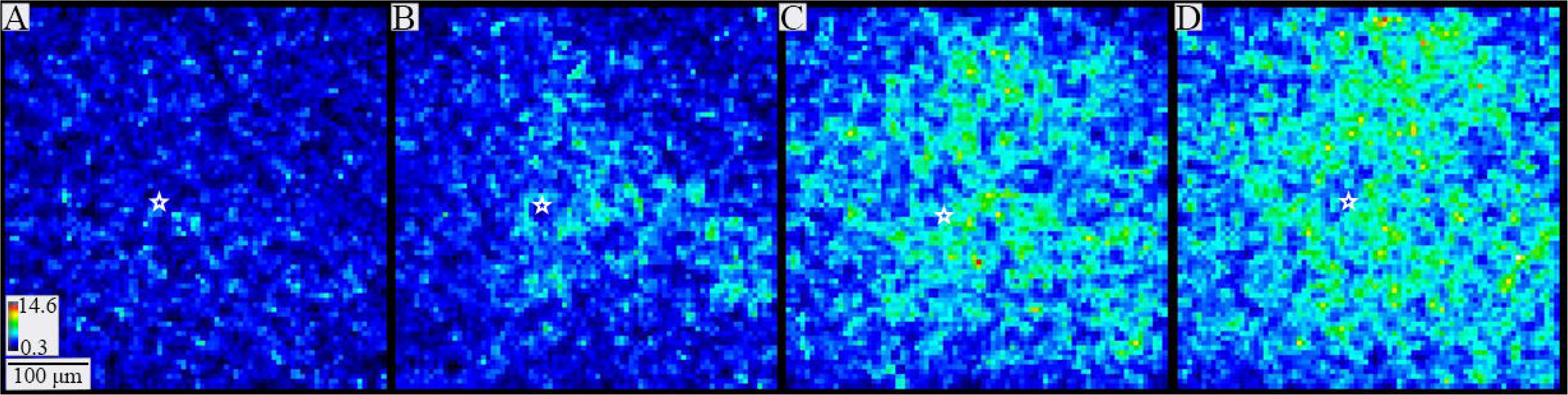
PV interneuron responses to different stimulation currents. Heatmaps of SNR following 10 μA (A), 20 μA (B), 60 μA (C), and 100 μA (D) stimulation. As current intensity increases, responses are visible through progressively larger areas. The stimulation site is marked with a white star in each panel. Heatmap scale in A also applies to B-D.

### Data processing, analysis, and velocity determination

Fluorescence divided by resting light intensity (ΔF/F) was passed through a nine-point binomial temporal filter and a spatial filter with σ = 1. A polynomial baseline correction was calculated using fluorescence outside of a 20 msec measurement window from 4 msec before to 16 msec after the stimulus. Signal-to-noise ratio (SNR) was calculated as the maximum amplitude divided by the standard deviation of noise in a 50-msec pre-stimulus window.

Signals in PV interneurons were used to track propagation through columns and layers by selecting contiguous groups of pixels as regions of interest (ROIs). A sequence of ROIs approximately 20 μm thick was automatically drawn along the hypothesized direction of propagation (illustrated in Fig. 3C), with widths spanning the barrel. Propagation distances used in the interlaminar velocity calculations were the distances from the stimulation site to each ROI center along the propagation trajectory parallel to the column edges. An analogous process was employed for intralaminar velocities between columns and within layers. Within each ROI, traces of fluorescence versus time were averaged and used to determine latency, defined as the time from stimulation to half-maximal response (illustration in Fig. 3D).

**Figure 3.**
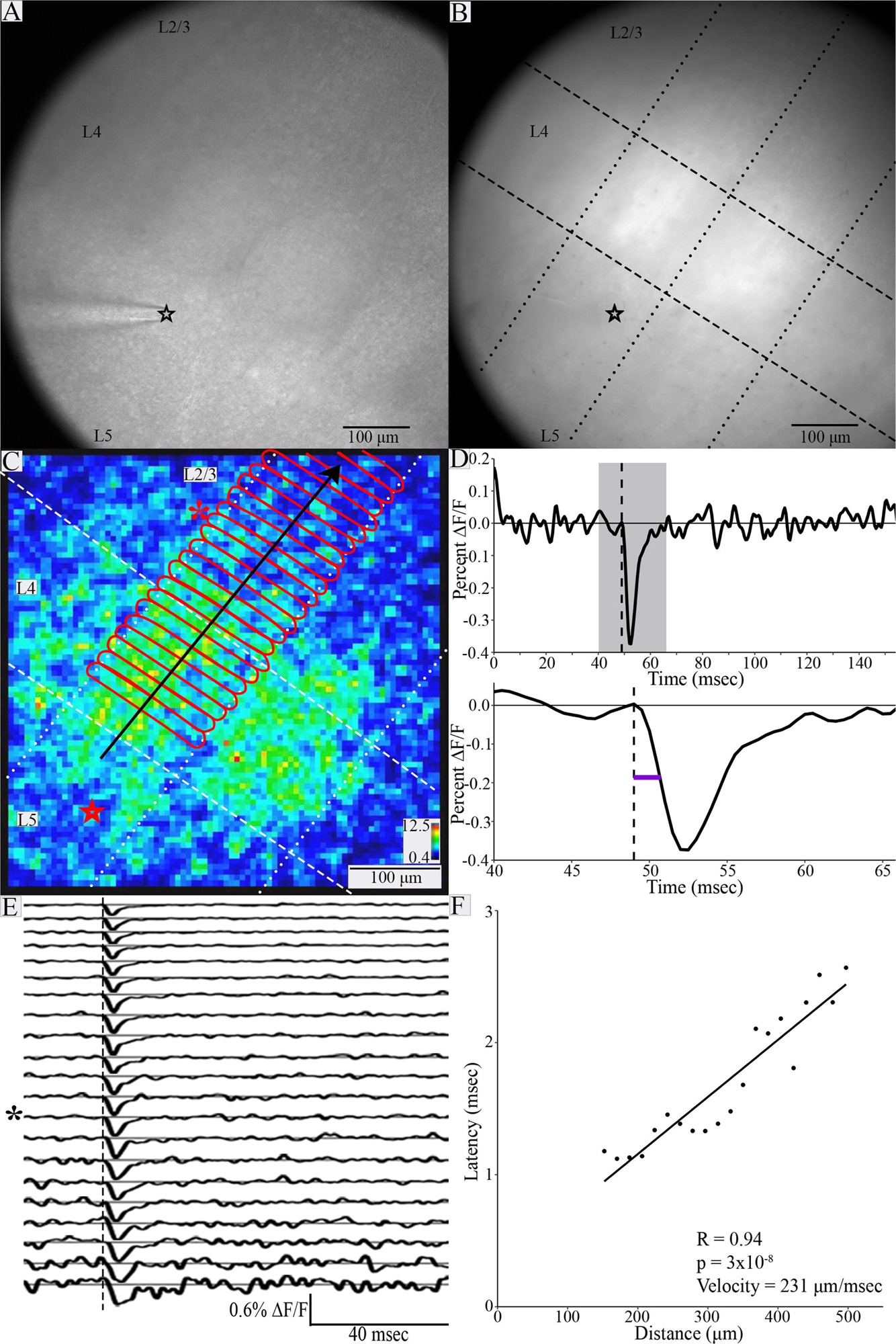
Spread of PV interneuron responses and determination of conduction velocity. Gradient contrast image (A) and fluorescence image (B) of a BC slice (both taken with the high-resolution Kiralux camera) with stimulating electrode in L5 (black star at the tip). Layer boundaries and barrels are visible in L4 in A. In B, layer boundaries are indicated with dashed lines, and column boundaries are indicated with dotted lines (in C as well). C. Heatmap of response SNR from the slice shown in A and B. Stimulation site indicated by red star (position differs slightly from A and B due to different cameras). Red outlines define ROIs approximately 20 μm thick spanning the stimulated column. Black arrow indicates direction of propagation. D. Top: Trace of fluorescence versus time for ROI indicated by asterisk in C and E. Gray shading indicates the 20 msec measurement window expanded below. Bottom: Latency (purple line) is the time from stimulation (dashed line) to half-maximal change in fluorescence of the rising phase. E. Traces of fluorescence versus time for 21 ROIs in sequence from bottom up (increasing distance from stimulation electrode). Dashed line indicates stimulation time, and asterisk indicates trace corresponding to the ROI marked with asterisk in C (thirteenth from bottom). Traces show PV interneuron responses with advancing latency. F. Latency plotted versus distance from the site of stimulation. The relationship between latency and distance was significant (R = 0.94, p = 3×10^-8^). The inverse of the slope gave an interlaminar conduction velocity of 231 μm/msec.

In addition to monosynaptic responses, stimulation often evoked disynaptic components and direct responses to electrical stimulation. Responses within 45 μm of the tip of the stimulating electrode or with latencies less than 1 msec after stimulation were assumed to be direct and were excluded. Additional direct and disynaptic responses were identified using sequences of SNR heatmaps, together with traces of fluorescence versus time. Only responses that were clearly monosynaptic were used for velocity determination. Conduction velocity was calculated as the inverse of the slope of a linear fit to plots of distance versus latency (illustrated in Fig. 3F), using plots with at least 7 points and with statistically significant correlations.

### Experimental design and statistical tests

Analysis included 31 slices from 13 animals (6 male and 7 female). Relationships between latency and distance were evaluated with linear regression, and p-values were corrected for multiple tests using the false discovery rate. To determine the appropriate statistical tests, normality was evaluated with Shapiro-Wilk tests, and differences in group variances were evaluated with Levene’s tests. Conduction velocity was normally distributed (W = 0.940, p = 0.084). Variance did not differ significantly for male and female animals (F(1, 29) = 0.001, p = 0.974) or for inter- and intralaminar velocities (F(1, 29) = 1.721, p = 0.200). It also did not differ for interlaminar conduction velocity from L2/3 to L4 and L5 (L2/3 → L5), interlaminar conduction velocity from L4 to L2/3 (L4 → L2/3) or L4 to L5 (L4 → L5), interlaminar conduction velocity from L5 to L2/3 or L4 (L5 → L2/3), intralaminar L2/3 conduction velocity, or intralaminar L4 conduction velocity (F(4, 26) = 1.437, p = 0.250). Therefore, interlaminar versus intralaminar velocities, and velocities for male versus female animals were compared with *t*-tests. Conduction velocity for different trajectories was compared with ANOVA. Conduction velocity did not differ significantly between sexes (t(28.898) = -1.437, p = 0.161).

### Code accessibility

Custom software, R code, and Python code available on request.

## Results

### PV interneuron responses to stimulation in L2/3, L4, and L5

We crossed PV-Cre driver mice with hVOS-Cre reporter mice to generate double transgenics PV-Cre;hVOS. We have previously shown that in the brain of PV-Cre;hVOS mice, hVOS probe is expressed in 83% of PV interneurons with 99.2% specificity (Bayguinov et al., 2017). Fig. 1A-C presents gradient contrast images of BC slices from PV-Cre;hVOS mice, and Fig. 1D-F presents the fluorescence images of these slices (note that the images were taken with different cameras, so the fields of view do not align precisely). Cortical layers were identified by cell density and cell size (Woolsey and Van der Loos, 1970; Feldmeyer, 2012) using gradient contrast (Fig. 1A-1C) and fluorescence (Fig. 1D-1F) images, and boundaries between layers are marked with dashed lines (Figs. 1D-1L). Fields of view generally contained L2/3 through L5. Barrels were separated by faint “hollows” (Woolsey and Van der Loos, 1970; Feldmeyer, 2012) as well as stronger fluorescence in L4, and boundaries between barrel columns are marked by dotted lines (Figs. 1D-1L). We assessed the velocity of spread of PV interneuron responses following stimulation in L2/3, L4, and L5. This elicited fluorescence changes associated with the depolarization of PV interneurons. Responses were seen throughout a slice, and the distributions of these responses are illustrated in peak amplitude (Figs. 1G-I) and SNR heatmaps (Figs. 1J-1L). Warmer colors correspond to larger depolarizations. Compared to the SNR heatmaps, amplitude heatmaps are less clear due to spatial variations in noise level within the field of view. We therefore used SNR heatmaps for the remainder of this paper. Traces of fluorescence versus time from different locations (indicated by number and color) reveal corresponding variations in the magnitude of stimulus-evoked PV interneuron depolarization, where fluorescence decreases as voltage moves DPA toward the fluorescent protein of the hVOS probe (Fig. 1M-O). Dark blue regions of the heatmaps indicate the absence of responsive PV interneurons, and traces from those locations show no discernable stimulus-evoked fluorescence changes (traces 5 in Figs. 1M-1O).

Increasing the stimulation current elicited responses in more cells over greater distances. SNR heatmaps (Fig. 2) revealed that 10 μA stimulation rarely elicited detectable responses (Fig. 2A). Increasing the stimulus current to 20 μA (Fig. 2B), 60 μA (Fig. 2C), and 100 μA (Fig. 2D) depolarized more neurons over larger areas. The observation that stimulation as weak as 20 µA elicits responses hundreds of micrometers from the stimulation electrode indicates that action potentials in axons propagate throughout the slice, to elicit synaptic responses which have been shown to depend on glutamate receptor activations (Canales et al., 2022). Response patterns varied but often included interlaminar and intralaminar responses across multiple layers and columns. These excitatory responses propagated away from the stimulating electrode and extended to the edges of the field of view within roughly 10 msec. The time course of this spread was used to measure conduction velocity.

### Quantification of conduction velocity

To determine conduction velocity, latency was plotted versus distance. Following identification of barrel and layer boundaries in gradient contrast and fluorescence images (Fig. 3A-B), ROIs were delineated within the stimulated column (for interlaminar velocity) or within the stimulated layer (for intralaminar velocity) (Fig. 3C; see Methods). Stimulus-induced fluorescence changes reported the roughly synchronous depolarization of PV interneurons within each ROI (Fig. 3E), and latency was measured as the time from stimulation to half-maximal change in fluorescence (Fig. 3D). As stated in Methods, plots of latency versus distance were well fitted by lines and the conduction velocity was taken as the inverse of the slope (Fig. 3F). Despite showing PV interneuron response propagation, in 15 plots latency and distance were not significantly correlated. In 7 of these 15 slices, estimated conduction velocity was faster than the highest velocity obtained from slices with significant correlations. In these cases, the spread within the field of view may have been too fast to detect differences in latency, as latencies in these plots varied by only 0.65 ± 0.16 msec (mean ± SD), and average root-mean-square error in latency was 0.185. In the remainder of this work, we will focus on conduction velocity measurements with significant distance-latency correlations and recognize that this biases our analysis uniformly toward slower conduction.

### Interlaminar and intralaminar conduction velocity

We next assessed conduction velocity for different propagation trajectories. Inter- and intralaminar conduction velocities were determined for stimulation in L2/3, L4, or L5. For L2/3 → L5 conduction velocity, we identified slices with clear propagation of PV interneuron responses within the stimulated column. Of 5 slices with 7 or more ROIs containing responses, 2 had significant correlations between latency and distance. In both slices, responses spanned L2/3 → L5, and their conduction velocity was 366 ± 9 μm/msec (mean ± SD, N = 2). Figure 4A displays three sequential SNR heatmaps at 1 msec intervals. Note that these are snapshots at specific times rather than peak amplitude heatmaps as in Figs. 1-3. This sequence of snapshots illustrates L2/3 → L5 conduction and Fig. 4B displays the associated plot of latency versus distance, yielding a velocity of 360 µm/sec.

**Figure 4.**
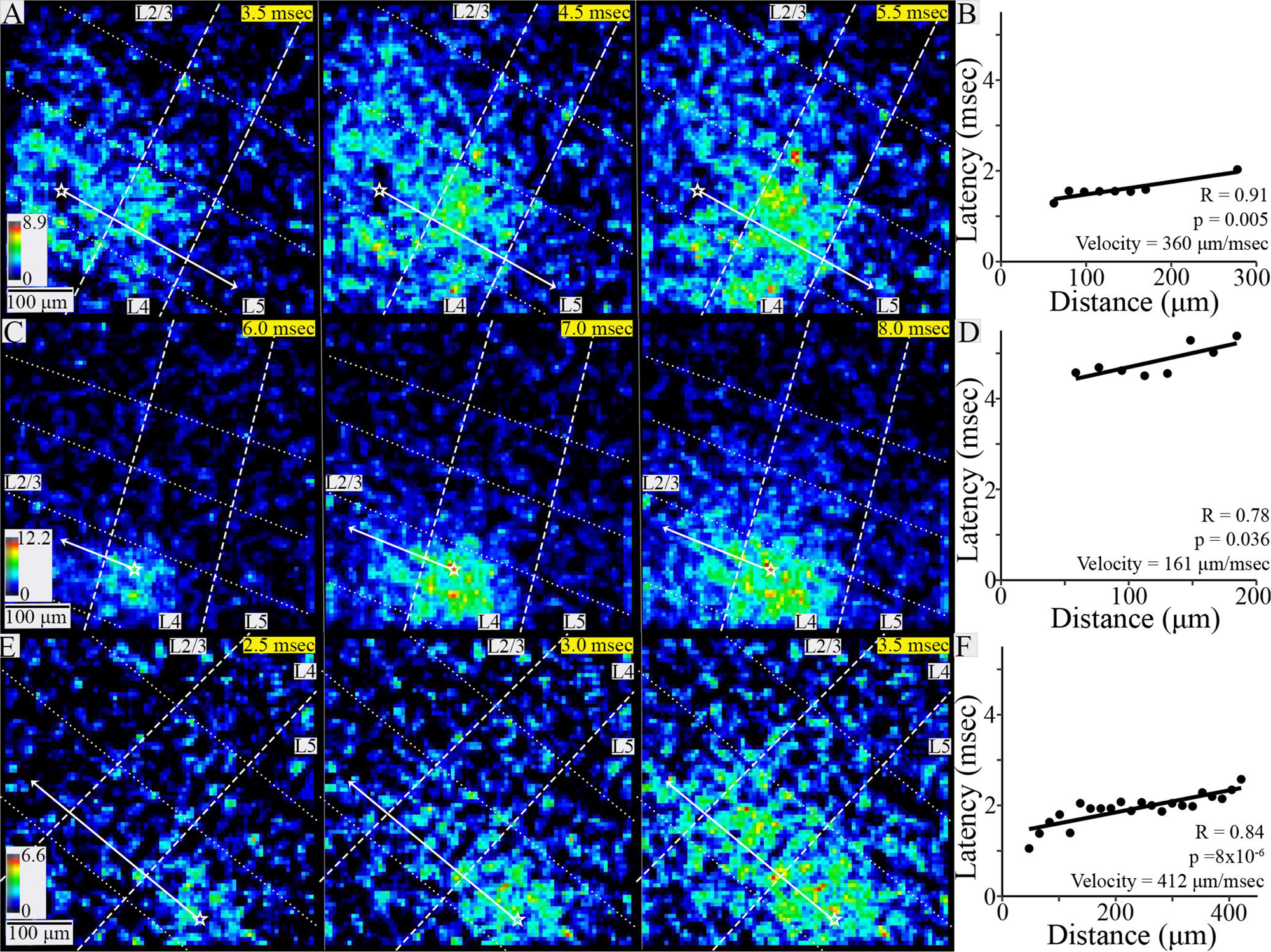
Interlaminar conduction. A. SNR heatmaps at 1-msec intervals showing the spread of PV interneuron responses to stimulation in L2/3. B. Latency versus distance within the stimulated column in the slice shown in A (R = 0.91, p = 0.005) yielded a conduction velocity of 360 μm/msec. C. SNR heatmaps at 1-msec intervals showing PV interneuron responses to stimulation in L4. D. Latency versus distance within the stimulated column in the slice shown in C (R = 0.78, p = 0.036) yielded a conduction velocity of 161 μm/msec. E. SNR heatmaps at 0.5-msec intervals showing PV interneurons responses to stimulation in L5. F. Latency versus distance within the stimulated column in the slice shown in E (R = 0.84, p = 8×10^-6^) yielded a conduction velocity of 412 μm/msec. In all SNR heatmaps, white stars mark stimulation site and solid white arrows show propagation trajectories used for velocity determination; dashed lines show layer boundaries, and dotted lines show column boundaries. Time after stimulation is shown in the upper right corner of each heatmap on yellow background.

For interlaminar responses to L4 stimulation, 3 of 6 slices yielded plots with significant correlations. L4 → L2/3 interlaminar conduction velocity was 223 ± 87 μm/msec (mean ± SD, N =2, example in Fig. 4C-D), and L4 → L5 interlaminar conduction velocity was 414 μm/msec. Overall, combined L4 → L2/3 and L4 → L5 conduction velocity was 286 ± 127 μm/msec (mean ± SD, N = 3).

In response to L5 stimulation, 16 of 18 slices produced significant correlations, and responses spread to L2/3 in all cases. L5 → L2/3 conduction velocity was 296 ± 122 μm/msec (mean ± SD, N = 16, example in Fig. 4E-F). The greater number of interlaminar velocity measurements compared to other trajectories was primarily due to less interference from disynaptic responses.

Of the 9 slices showing L2/3 intralaminar conduction, latency and distance were significantly correlated in 6, yielding a conduction velocity of 200 ± 93 μm/msec (mean ± SD, N = 6). Responses were restricted to the stimulated and immediately adjacent columns (Fig. 5A-B) in all but one slice. In that slice responses extended to one additional column. Of 6 slices exhibiting L4 intralaminar conduction, latency and distance were significantly correlated in 4, yielding a conduction velocity of 142 ± 76 μm/msec (mean ± SD, N = 4). Responsive ROIs spanned the home and neighboring columns in 2 slices (Fig. 5C-D) and extended to an additional adjacent column in 2 more slices. Finally, in the 2 slices exhibiting L5 intralaminar conduction, latency and distance were not correlated and no velocity was determined.

**Figure 5.**
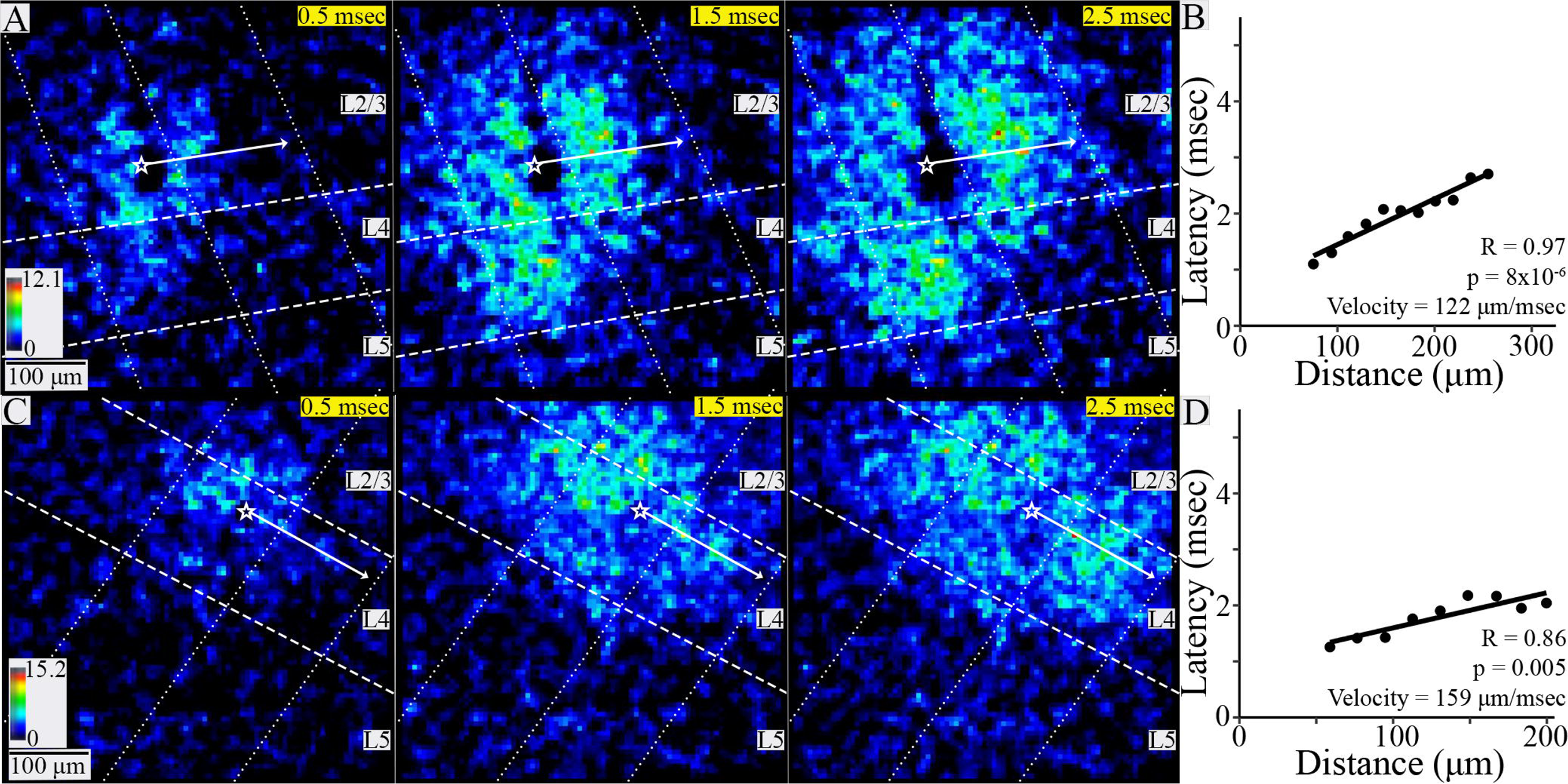
Intralaminar conduction. A. SNR heatmaps at 1-msec intervals showing the spread of PV interneuron responses within L2/3 to stimulation in that layer. Responses spread within the stimulated column and to a neighboring column. B. Latency versus distance for the slice shown in A (R = 0.97, p = 8×10^-6^) yielded a conduction velocity of 112 μm/msec. C. SNR heatmaps at 1-msec intervals showing PV interneuron responses within L4 to stimulation in that layer. D. Latency versus distance for the slice shown in C (R = 0.86, p = 0.005) yielded a conduction velocity of 159 μm/msec. In all heatmaps, white stars mark stimulation site and white arrows show PV interneuron response propagation trajectories; dashed lines show layer boundaries, and dotted lines show column boundaries. Time after stimulation shown in upper right corner of each heatmap on yellow background.

Combining measurements in 21 slices from 11 animals, average interlaminar conduction velocity to stimulation in L2/3, L4, and L5 was 302 ± 115 μm/msec (mean ± SD). L2/3 and L4 intralaminar conduction velocity in 10 slices from 7 animals was 177 ± 87 μm/msec (mean ± SD). Interlaminar conduction velocity was therefore about 71% faster than intralaminar conduction velocity (Fig. 6A), and this difference was significant (t(22.996) = 3.344, p = 0.003, Welch’s two-sample t-test). L4 → L2/3 or L5 and L5 → L2/3 conduction velocity were not significantly different (t(2.740) = -0.124, p = 0.910, Welch’s two-sample t-test, Fig. 6B). L2/3 and L4 intralaminar conduction velocity also were not significantly different (t(7.506) = 1.086, p = 0.311, Welch’s two-sample t-test, Fig. 6B). One slice produced both interlaminar and intralaminar conduction. In this slice, L4 → L5 conduction (414 μm/msec) was more than twice as fast as intralaminar L4 conduction (159 μm/msec), in keeping with the trend of more rapid interlaminar conduction. In summary, we measured conduction velocity of excitatory axons targeting PV interneurons across multiple trajectories, and found that interlaminar conduction is significantly faster than intralaminar conduction.

**Figure 6.**
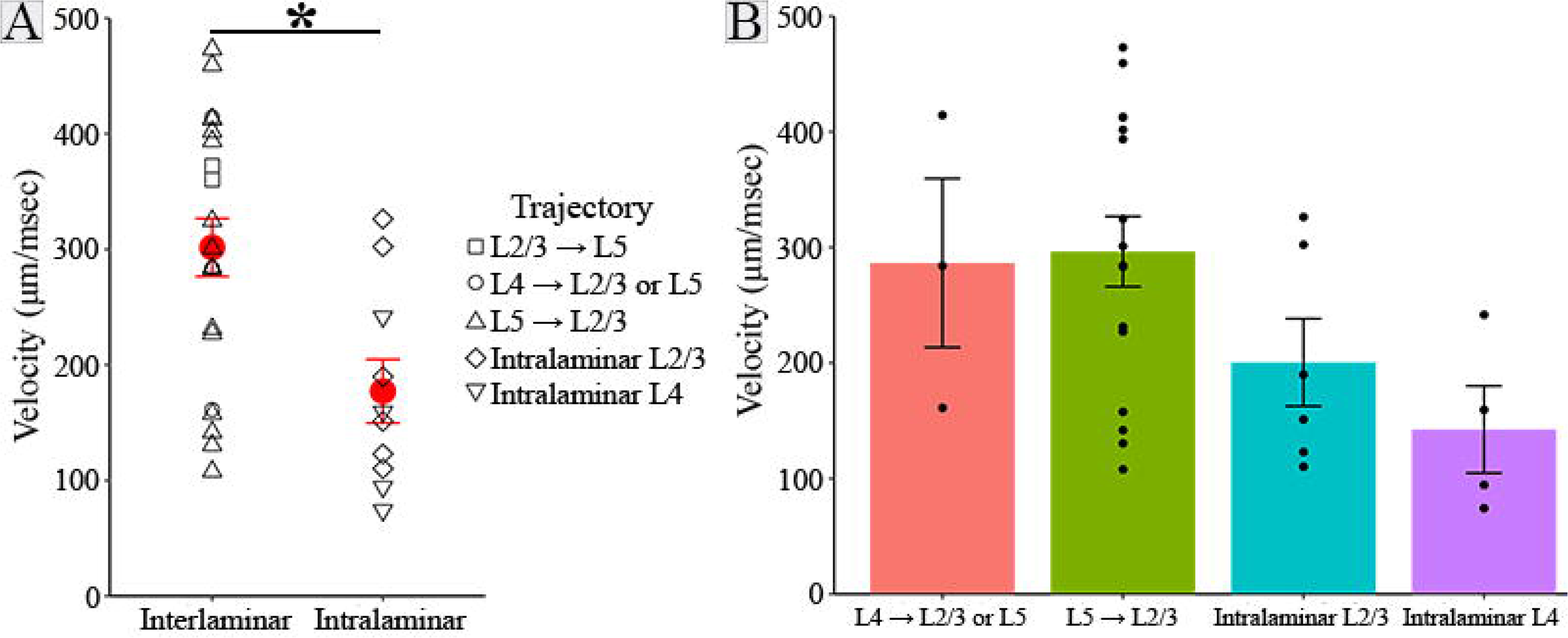
Interlaminar and intralaminar conduction velocity. A. Interlaminar velocity (302 ± 115 μm/msec, 21 slices) was approximately 71% faster than intralaminar velocity (177 ± 87 μm/msec, 10 slices; t(22.996) = 3.344, p = 0.003, Welch’s two-sample t-test). Symbol shapes indicate velocity trajectory (in legend), and group mean ± SE are shown in red. B. L4 → L2/3 or L5 (286 ± 127 μm/msec, 3 slices), L5 → L2/3 (296 ± 122 μm/msec, 16 slices), L2/3 intralaminar (200 ± 93 μm/msec, 6 slices) and L4 intralaminar (142 ± 76 μm/msec, 4 slices) velocity. Error bars indicate SE.

## Discussion

In this study we used voltage imaging from PV interneurons to investigate the spread of excitation through interlaminar and intralaminar circuits in the BC. The PV interneuron response patterns recapitulate known circuitry across both canonical and non-canonical pathways. Canonically, thalamic input enters the cortex through L4, and is then relayed to L2/3 (Xu and Callaway, 2009; Staiger and Petersen, 2021). L4 provides strong excitatory drive to L2/3 PV interneurons (Helmstaedter et al., 2008; Adesnik et al., 2012), and we determined L4 → L2/3 conduction velocity to be 223 ± 87 μm/msec (mean ± SD). While both basket cells and chandelier cells, two morphologically distinct PV interneuron subtypes, are present in L2/3, L4 neurons do not target L2/3 chandelier cells (Xu and Callaway, 2009). This velocity therefore reflects conduction along excitatory cell axons specifically targeting L2/3 basket cells. In addition to the canonical L4 → L2/3 interlaminar path, spiny stellate cells in L4 are also known to project to PV interneurons in L5 (Pluta et al., 2015). We determined L4 → L5 conduction velocity to be 414 μm/msec based on one slice. It is interesting that the L4 → L5 conduction velocity measured here (414 μm/msec) was faster than both of our L4 → L2/3 conduction velocity values (161 μm/msec and 284 μm/msec).

Continuing along the canonical cortical circuit, L2/3 pyramidal cells project to L5 excitatory and inhibitory cells, including PV interneurons (Lourenco et al., 2020). Stimulating in L2/3 elicited responses in PV interneurons in both L4 and L5. With a velocity of 366 ± 9 μm/msec (N = 2), L2/3 → L5 conduction was the fastest and least variable velocity found in the present study. Finally, L5 stimulation produced responses in L4 and L2/3 PV interneurons with a velocity of 296 ± 122 μm/msec (mean ± SD, N = 16). L2/3 chandelier cells receive especially strong input from L5A (Xu and Callaway, 2009). Because we stimulated mainly in the upper portion of L5, this velocity may reflect conduction along this projection. Overall, the basic patterns of spread we see recapitulate the known functional connectivity between cortical layers in BC, and our work provides velocities for conduction along these projections.

Our findings are also consistent with intralaminar connectivity in BC. Previous anatomical findings have reported that L2/3 pyramidal cell axons extend horizontally into neighboring columns (Narayanan et al., 2015), and L2/3 stimulation in rat primary somatosensory cortex can elicit intralaminar responses up to 2 mm away (Telfeian and Connors, 2003). In L4, spiny stellate cell axons also sometimes cross into a neighboring barrel (Egger et al., 2008; Staiger and Petersen, 2021). We report L2/3 intralaminar (200 ± 93 μm/msec, mean ± SD, N = 6) and L4 intralaminar (142 ± 76 μm/msec, mean ± SD, N = 4) conduction velocity through these intralaminar circuits. Intralaminar PV interneuron responses remained within the stimulated column and a neighboring column in 7 experiments, but spread to an additional column in 3 experiments. This supports the concept of an “intracortical unit” spanning three barrel columns (Narayanan et al., 2015). Although L5 intralaminar axons can extend through up to two neighboring columns, we observed intralaminar L5 conduction in only 2 slices and were unable to measure any conduction velocities along this trajectory. We are thus imaging conduction along previously described intralaminar and interlaminar circuits within BC, and again providing velocities of propagation along these trajectories.

Our range of conduction velocities from 31 measurements (74 – 473 μm/msec) is broadly consistent with previous conduction velocity measurements along murine hippocampal, intracortical, and thalamocortical axons. This includes measurements with various methods analogous to ours using postsynaptic responses to track conduction, including antidromic activation (300 μm/msec, (Shlosberg et al., 2008)), microelectrode arrays (330 μm/msec, (Bakker et al., 2009)), and extracellular stimulation and patch clamp recordings (363 μm/msec, (Salami et al., 2003); 150-550 μm/msec, (Murakoshi et al., 1993); 400 μm/msec, (Telfeian and Connors, 2003); 200 μm/msec, (Helmstaedter et al., 2008)). Our velocity values are also consistent with previously published optical measurements directly from axons with voltage sensitive dye delivered to individual cells through a patch pipette (50-450 μm/msec, Popovic et al., 2011) or with axon-targeted hVOS measurements (94 and 228 μm/msec, Ma et al., 2017).

We report that interlaminar conduction velocity (302 ± 115 μm/msec, mean ± SD, N = 21) is 71% faster than intralaminar conduction velocity (177 ± 87 μm/msec, mean ± SD, N = 10). Because functions differ for interlaminar and intralaminar circuits, these differences in conduction velocity likely have functional implications. L4 → L2/3 conduction velocity impacts integration of intracortical and thalamic input in L2/3 (Kimura et al., 2010). Optogenetic inhibition of L4 excitatory cells broadens L5 excitatory cell sensory tuning curves by increasing responses to non-preferred stimuli, and this effect is likely mediated by L5 PV interneurons (Pluta et al., 2015). Stress decreases PV interneuron activity across L2-L5, leads to loss of dendritic spines on L5 pyramidal cells, and impairs texture discrimination. Stimulation of PV interneurons ameliorates these deficits (Chen et al., 2018). Finally, PV interneurons are hypothesized to decrease synchrony between L4 and L5 during active sensory periods (Jang et al., 2020), and small changes in conduction velocity can have large impacts on synchrony (Pajevic et al., 2014; Ivanov et al., 2019). Interlaminar conduction velocity may therefore impact functions such as texture discrimination, intracortical synchrony, sensory tuning, and integration of thalamic and intracortical input.

In contrast, timing of intralaminar input to PV interneurons from neighboring columns likely plays a role in multi-whisker integration. In L2/3, each barrel contains pinwheel shaped directional preference maps resembling pinwheel-shaped orientation selectivity maps in V1. Both putative pyramidal cells and inhibitory interneurons show directional preferences (Andermann and Moore, 2006). In visual cortex L2/3 intralaminar axons between columns might connect regions with similar preferred directions (Gilbert, 1992). L3 and L4 cells in columns corresponding to whiskers along the same arc have similar preferred whisker stimulation frequencies (Andermann et al., 2004), and L2/3 excitation spreads across columns related to whiskers along rows *before* arcs (Petersen et al., 2003). L2/3 and L4 intralaminar conduction velocity may therefore impact multi-whisker integration of frequency and direction preferences differentially based on input to whiskers within rows and arcs.

Overall, interlaminar conduction is approximately 71% faster than intralaminar conduction. Consequently, computations within a barrel column may occur more quickly than those across columns. Thus, multi-whisker integration, which depends on intralaminar communication between barrel columns, will occur more slowly than computations within columns related to intracortical synchrony, sensory tuning, and texture discrimination.

hVOS imaging offers a new and powerful approach to the study of conduction velocity, not only in the axons of defined cell types (Ma et al., 2017), but also in the axons defined by their targeted cell types. Although DPA increases membrane capacitance and thus slows propagation, this effect is small as increasing DPA from 2 to 4 μM reduced conduction velocity by only 15% in mossy fibers (Ma et al., 2017). Furthermore, this non-specific effect will reduce conduction velocity proportionally in different populations of axons. Voltage imaging has historically provided a powerful method for the measurement of axonal conduction velocity (Grinvald et al., 1981; Sakai et al., 1991; Popovic et al., 2011; Hamada et al., 2017), and genetically encoded voltage indicators have extended this approach through the addition of targeting specificity (Ma et al., 2019; Panzera and Hoppa, 2019). Conduction velocity can vary widely based on many factors, including the identity of the postsynaptic cell being targeted, and differences in velocity have functional implications (Kimura et al., 2010). Because it can be genetically targeted to specific cell populations (Bayguinov et al., 2017). hVOS offers a unique opportunity to study differences in conduction velocity along axons based on their postsynaptic targets. Targeting hVOS probes to different types of neurons thus has the potential to reveal additional forms of axon specialization adapted to different forms of neuronal computations.

## Acknowledgements

National Institutes of Health Grants NS127219 and NS093866 to M.B.J. and NS105200 X.Z. J.J was supported by T32GM130550. Thanks to Dr. Shane McMahon for methodological contributions.

## Notes

### Competing Interest Statement

The authors have declared no competing interest.

### Summary of Updates

Improved method of analysis

## References

Adesnik H (2018) Layer-specific excitation/inhibition balances during neuronal synchronization in the visual cortex. J Physiol-London 596:1639–1657.

Adesnik H, Bruns W, Taniguchi H, Huang ZJ, Scanziani M (2012) A neural circuit for spatial summation in visual cortex. Nature 490:226–231.

Allitt BJ, Alwis DS, Rajan R (2017) Laminar-specific encoding of texture elements in rat barrel cortex. J Physiol 595:7223–7247.

Andermann ML, Moore CI (2006) A somatotopic map of vibrissa motion direction within a barrel column. Nature Neuroscience 9:543–551.

Andermann ML, Ritt J, Neimark MA, Moore CI (2004) Neural Correlates of Vibrissa Resonance: Band-Pass and Somatotopic Representation of High-Frequency Stimuli. Neuron 42.

Ayaz A, Stauble A, Hamada M, Wulf MA, Saleem AB, Helmchen F (2019) Layer-specific integration of locomotion and sensory information in mouse barrel cortex. Nat Commun 10:2585.

Bakker R, Schubert D, Levels K, Bezgin G, Bojak I, Kotter R (2009) Classification of cortical microcircuits based on micro-electrode-array data from slices of rat barrel cortex. Neural Netw 22:1159–1168.

Bayguinov PO, Ma YH, Gao Y, Zhao XY, Jackson MB (2017) Imaging Voltage in Genetically Defined Neuronal Subpopulations with a Cre Recombinase-Targeted Hybrid Voltage Sensor. Journal of Neuroscience 37:9305–9319.

Bradley J, Luo R, Otis TS, DiGregorio DA (2009) Submillisecond Optical Reporting of Membrane Potential In Situ Using a Neuronal Tracer Dye. Journal of Neuroscience 29:9197–9209.

Canales A, Scheuer KS, Zhao X, Jackson MB (2022) Unitary synaptic responses of parvalbumin interneurons evoked by excitatory neurons in the mouse barrel cortex. Cereb Cortex.

Cardin JA (2018) Inhibitory Interneurons Regulate Temporal Precision and Correlations in Cortical Circuits. Trends in Neurosciences 41:689–700.

Chanda B, Blunck R, Faria LC, Schweizer FE, Mody I, Bezanilla F (2005) A hybrid approach to measuring electrical activity in genetically specified neurons. Nat Neurosci 8:1619–1626.

Chang PY (2006) Heterogeneous spatial patterns of long-term potentiation in hippocampal slices. In: Biophysics, p 144. Madison, WI: University of Wisconsin-Madison.

Chen CC, Lu J, Yang R, Ding JB, Zuo Y (2018) Selective activation of parvalbumin interneurons prevents stress-induced synapse loss and perceptual defects. Mol Psychiatry 23:1614–1625.

Egger V, Nevian T, Bruno RM (2008) Subcolumnar dendritic and axonal organization of spiny stellate and star pyramid neurons within a barrel in rat somatosensory cortex. Cereb Cortex 18:876–889.

Feldmeyer D (2012) Excitatory neuronal connectivity in the barrel cortex. Front Neuroanat 6:24.

Ferguson BR, Gao WJ (2018) PV Interneurons: Critical Regulators of E/I Balance for Prefrontal Cortex-Dependent Behavior and Psychiatric Disorders. Front Neural Circuits 12:37.

Galarreta M, Hestrin S (2002) Electrical and chemical synapses among parvalbumin fast-spiking GABAergic interneurons in adult mouse neocortex. P Natl Acad Sci USA 99:12438–12443.

Ghitani N, Bayguinov PO, Ma YH, Jackson MB (2015) Single-trial imaging of spikes and synaptic potentials in single neurons in brain slices with genetically encoded hybrid voltage sensor. Journal of Neurophysiology 113:1249–1259.

Gilbert CD (1992) Horizontal Integration and Cortical Dynamics. Neuron 9:1–13.

Gonzalez-Burgos G, Cho RY, Lewis DA (2015) Alterations in cortical network oscillations and parvalbumin neurons in schizophrenia. Biol Psychiatry 77:1031–1040.

Grinvald A, Ross WN, Farber I (1981) Simultaneous optical measurements of electrical activity from multiple sites on processes of cultured neurons. Proc Natl Acad Sci U S A 78:3245–3249.

Hamada MS, Popovic MA, Kole MH (2017) Loss of Saltation and Presynaptic Action Potential Failure in Demyelinated Axons. Front Cell Neurosci 11:45.

Helmstaedter M, Staiger JF, Sakmann B, Feldmeyer D (2008) Efficient recruitment of layer 2/3 interneurons by layer 4 input in single columns of rat somatosensory cortex. J Neurosci 28:8273–8284.

Isett BR, Feasel SH, Lane MA, Feldman DE (2018) Slip-Based Coding of Local Shape and Texture in Mouse S1. Neuron 97:418–433 e415.

Ivanov VA, Polykretis IE, Michmizos KP (2019) Axonal Conduction Velocity Impacts Neuronal Network Oscillations. 2019 Ieee Embs International Conference on Biomedical & Health Informatics (Bhi).

Jang HJ, Chung HW, Rowland JM, Richards BA, Kohl MM, Kwag JY (2020) Distinct roles of parvalbumin and somatostatin interneurons in gating the synchronization of spike times in the neocortex. Sci Adv 6.

Kimura F, Itami C, Ikezoe K, Tamura H, Fujita I, Yanagawa Y, Obata K, Ohshima M (2010) Fast activation of feedforward inhibitory neurons from thalamic input and its relevance to the regulation of spike sequences in the barrel cortex. J Physiol-London 588:2769–2787.

Lauber E, Filice F, Schwaller B (2018) Parvalbumin neurons as a hub in autism spectrum disorders. J Neurosci Res 96:360–361.

Lefort S, Tomm C, Floyd Sarria JC, Petersen CC (2009) The excitatory neuronal network of the C2 barrel column in mouse primary somatosensory cortex. Neuron 61:301–316.

Lenschow C, Brecht M (2015) Barrel cortex membrane potential dynamics in social touch. Neuron 85:718–725.

Liu TY, Hsieh JC, Chen YS, Tu PC, Su TP, Chen LF (2012) Different patterns of abnormal gamma oscillatory activity in unipolar and bipolar disorder patients during an implicit emotion task. Neuropsychologia 50:1514–1520.

Lourenco J, De Stasi AM, Deleuze C, Bigot M, Pazienti A, Aguirre A, Giugliano M, Ostojic S, Bacci A (2020) Modulation of Coordinated Activity across Cortical Layers by Plasticity of Inhibitory Synapses. Cell Rep 30:630–641 e635.

Ma Y, Bayguinov PO, Jackson MB (2017) Action Potential Dynamics in Fine Axons Probed with an Axonally Targeted Optical Voltage Sensor. eNeuro 4.

Ma YH, Bayguinov PO, Jackson MB (2019) Optical studies of action potential dynamics with hVOS probes. Curr Opin Biomed Eng 12:51–58.

Murakoshi T, Guo JZ, Ichinose T (1993) Electrophysiological identification of horizontal synaptic connections in rat visual cortex in vitro. Neurosci Lett 163:211–214.

Narayanan RT, Egger R, Johnson AS, Mansvelder HD, Sakmann B, de Kock CP, Oberlaender M (2015) Beyond Columnar Organization: Cell Type- and Target Layer-Specific Principles of Horizontal Axon Projection Patterns in Rat Vibrissal Cortex. Cereb Cortex 25:4450–4468.

O’Connor DH, Peron SP, Huber D, Svoboda K (2010) Neural activity in barrel cortex underlying vibrissa-based object localization in mice. Neuron 67:1048–1061.

Pajevic S, Basser PJ, Fields RD (2014) Role of myelin plasticity in oscillations and synchrony of neuronal activity. Neuroscience 276:135–147.

Panzera LC, Hoppa MB (2019) Genetically Encoded Voltage Indicators Are Illuminating Subcellular Physiology of the Axon. Front Cell Neurosci 13:52.

Petersen CC, Grinvald A, Sakmann B (2003) Spatiotemporal dynamics of sensory responses in layer 2/3 of rat barrel cortex measured in vivo by voltage-sensitive dye imaging combined with whole-cell voltage recordings and neuron reconstructions. J Neurosci 23:1298–1309.

Pluta S, Naka A, Veit J, Telian G, Yao L, Hakim R, Taylor D, Adesnik H (2015) A direct translaminar inhibitory circuit tunes cortical output. Nat Neurosci 18:1631–1640.

Popovic MA, Foust AJ, McCormick DA, Zecevic D (2011) The spatio-temporal characteristics of action potential initiation in layer 5 pyramidal neurons: a voltage imaging study. J Physiol 589:4167–4187.

Pouille F, Scanziani M (2001) Enforcement of temporal fidelity in pyramidal cells by somatic feed-forward inhibition. Science 293:1159–1163.

Sakai T, Komuro H, Katoh Y, Sasaki H, Momose-Sato Y, Kamino K (1991) Optical determination of impulse conduction velocity during development of embryonic chick cervical vagus nerve bundles. J Physiol 439:361–381.

Salami M, Kimura F, Tsumoto T (2003) Postnatal changes of conduction velocity of the fibers in and out of the mouse barrel cortex. Iran Biomed J 7.

Shlosberg D, Abu-Ghanem Y, Amitai Y (2008) Comparative properties of excitatory and inhibitory inter-laminar neocortical axons. Neuroscience 155:366–373.

Staiger JF, Petersen CCH (2021) Neuronal Circuits in Barrel Cortex for Whisker Sensory Perception. Physiol Rev 101:353–415.

Telfeian AE, Connors BW (2003) Widely integrative properties of layer 5 pyramidal cells support a role for processing of extralaminar synaptic inputs in rat neocortex. Neurosci Lett 343:121–124.

Vecchia D, Beltramo R, Vallone F, Chereau R, Forli A, Molano-Mazon M, Bawa T, Binini N, Moretti C, Holtmaat A, Panzeri S, Fellin T (2020) Temporal Sharpening of Sensory Responses by Layer V in the Mouse Primary Somatosensory Cortex. Curr Biol 30:1589–1599 e1510.

Wang D, Zhang Z, Chanda B, Jackson MB (2010) Improved probes for hybrid voltage sensor imaging. Biophys J 99:2355–2365.

Woolsey TA, Van der Loos H (1970) The structural organization of layer IV in the somatosensory region (S1) of mouse cerebral cortex. Brain Res 17.

Xu X, Callaway EM (2009) Laminar specificity of functional input to distinct types of inhibitory cortical neurons. J Neurosci 29:70–85.

Xu XM, Olivas ND, Ikrar T, Peng T, Holmes TC, Nie Q, Shi YL (2016) Primary visual cortex shows laminar-specific and balanced circuit organization of excitatory and inhibitory synaptic connectivity. J Physiol-London 594:1891–1910.

Yeganeh F, Knauer B, Guimaraes Backhaus R, Yang JW, Stroh A, Luhmann HJ, Stuttgen MC (2022) Effects of optogenetic inhibition of a small fraction of parvalbumin-positive interneurons on the representation of sensory stimuli in mouse barrel cortex. Sci Rep 12:19419.

